## Supplementary Figures and Tables for "Staufen2 mediated RNA recognition and localization requires combinatorial action of multiple domains"

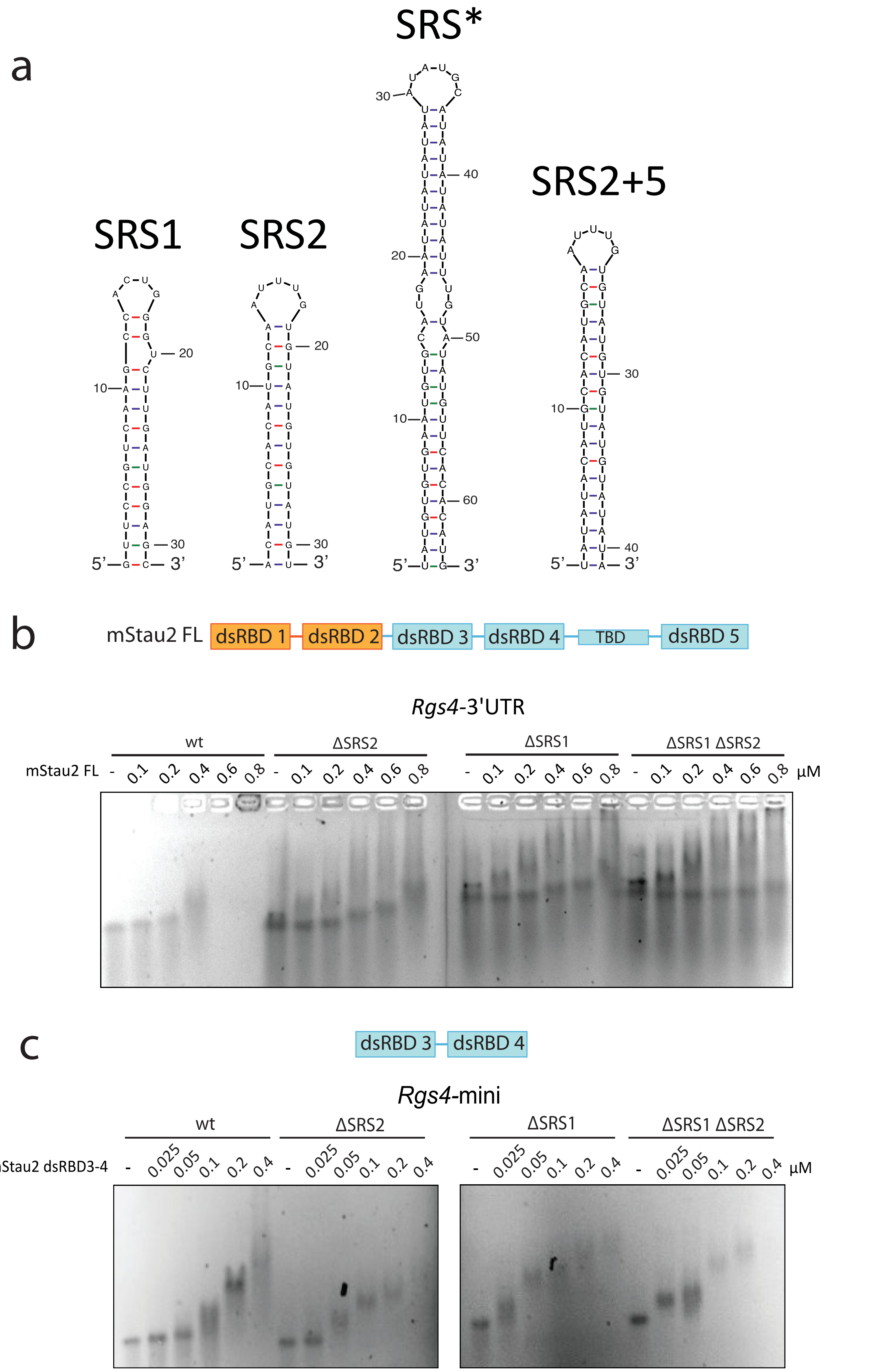


**Supplementary Figure 1: EMSAs with mStau2 and *Rgs4* 3'UTR RNAs. a** Schematic representation of the RNAs used in this study. SRS1 and SRS2 are predicted Staufen-recognized structures (SRS) in the *Rgs4* 3'UTR[1]. SRS* is the most stably predicted secondary structure in the *Rgs4* 3'UTR. SRS2+5 is an elongated version of SRS2. **b** mStau2 full-length binds wild type *Rgs4* 3'UTR and SRS deletion mutants with similar apparent affinities in the nanomolar concentration range. **c** mStau2 dsRBD3-4 binds *Rgs4*-mini wild type and SRS deletion mutants with similar apparent affinities in the nanomolar concentration range. Complexes were resolved in 1.5 % agarose gels and imaged via GelRed staining and UV imaging. Source data are provided as a Source Data file.

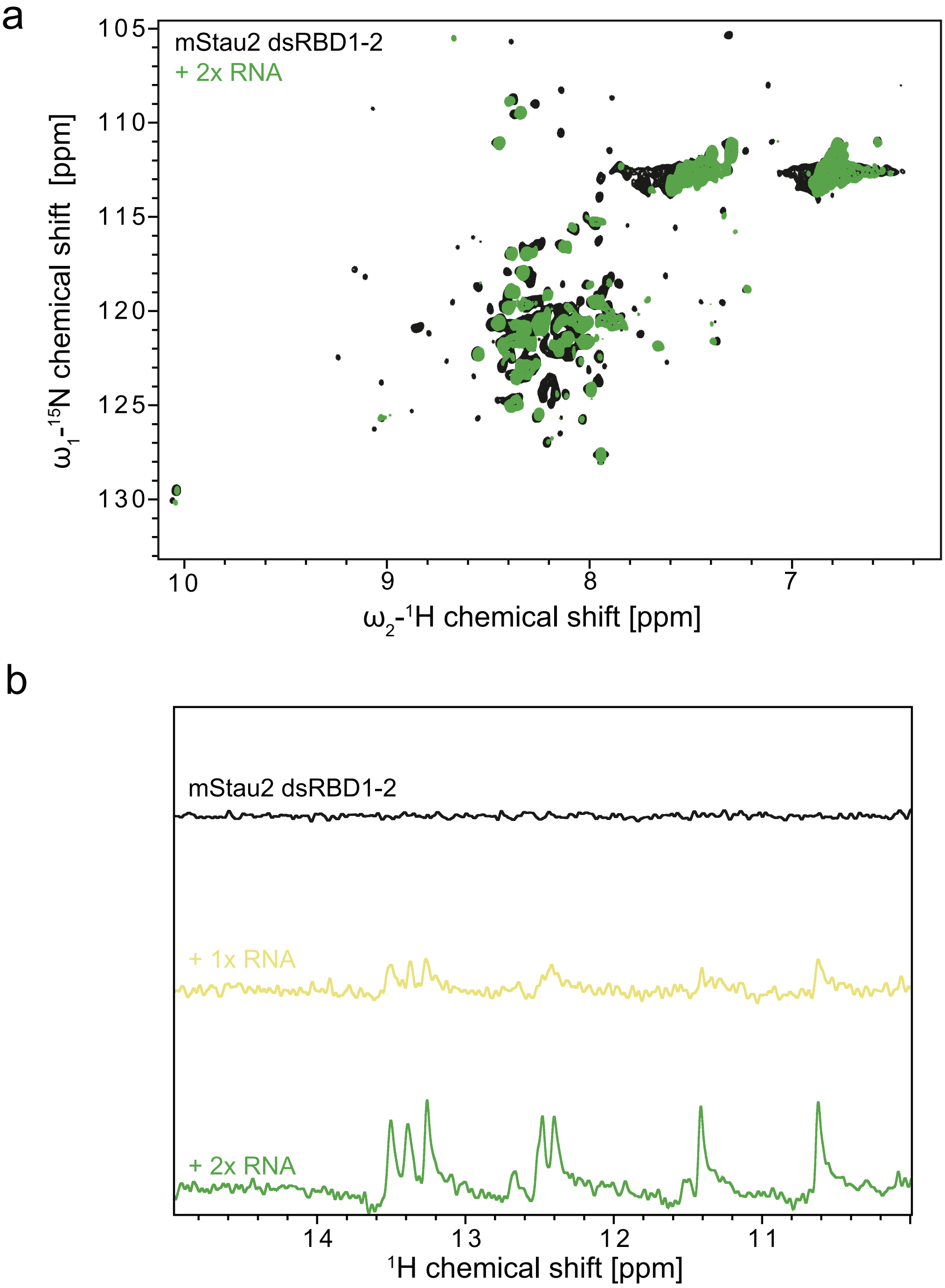


**Supplementary Figure 2: NMR titration experiments of Stau2 dsRBD1-2 with SRS2 RNA. a** Overlay of $^{1}H,^{15}N$-HSQC spectra of dsRBD1-2 in absence and presence of 2x excess SRS2 RNA. Resonance shifts and line broadening of several signals are observed. **b** Comparison of 1D imino traces of SRS2 RNA at different stoichiometric ratios with dsRBD1-2.

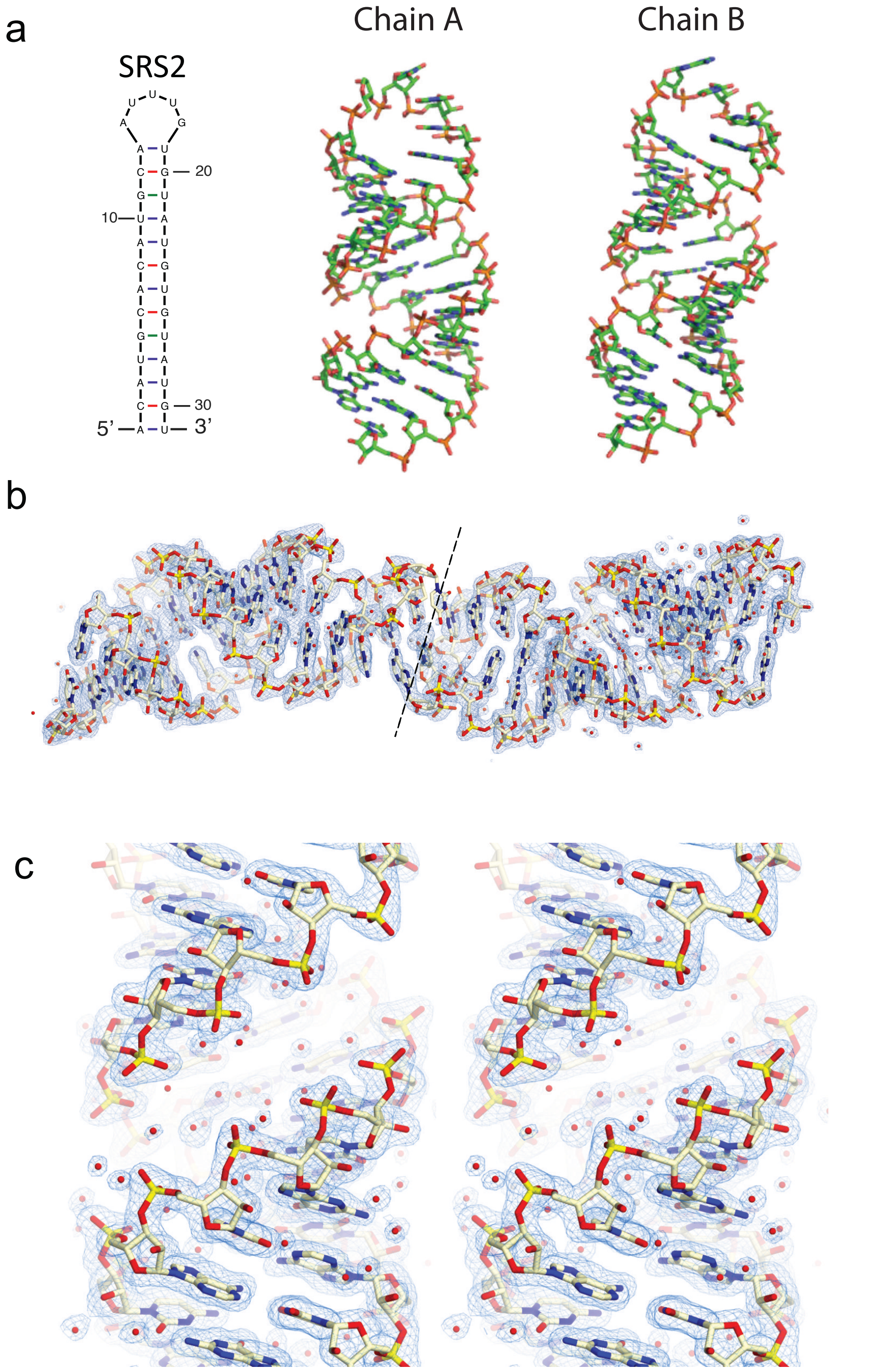


**Supplementary Figure 3: Crystal structure of the isolated *Rgs4* SRS2 RNA stem-loop.** **a** Schematic drawing of SRS2 RNA and the two molecules chain A and chain B contained in the asymmetric unit of the crystal lattice. Both molecules adopt the typical RNA A-form and form the expected stem-loop. Both chains differ slightly in their loop-regions, which appear to be disordered. **b** (2F(o)-F(c)) electron density map of the two RNA molecules in the asymmetric unit at 1σ contour. Whereas the density map in the stem region of the RNA is very well defined, it is rather poor in the area of the two loops, indicating disordered loop regions. **c** Stereo image of a fragment of the SRS2 stem showing the (2F(o)-F(c)) electron density map at 1σ contour.

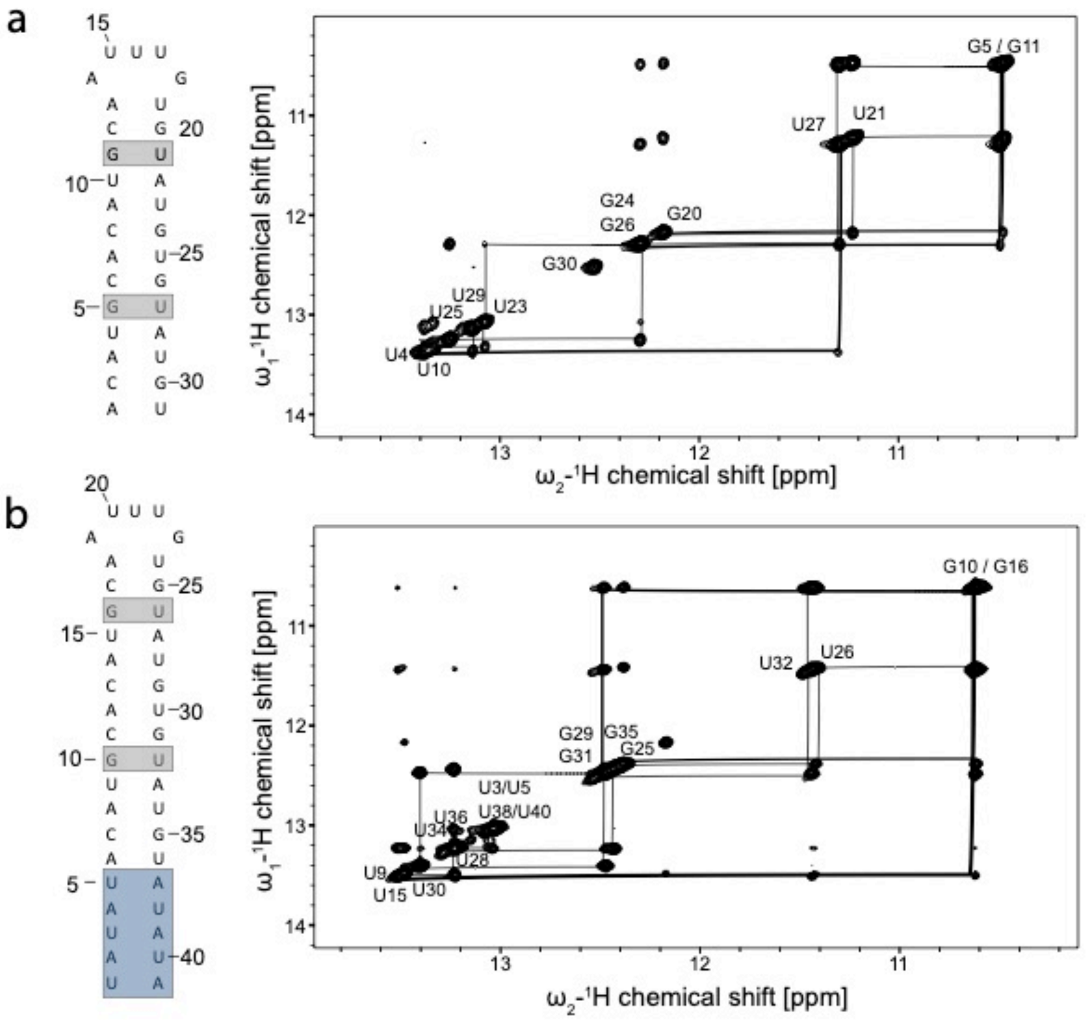


**Supplementary Figure 4: RNA assignment of imino groups based on $^1H$, $^1H$- NOESY spectra of a SRS2 and b SRS2+5 RNA.**

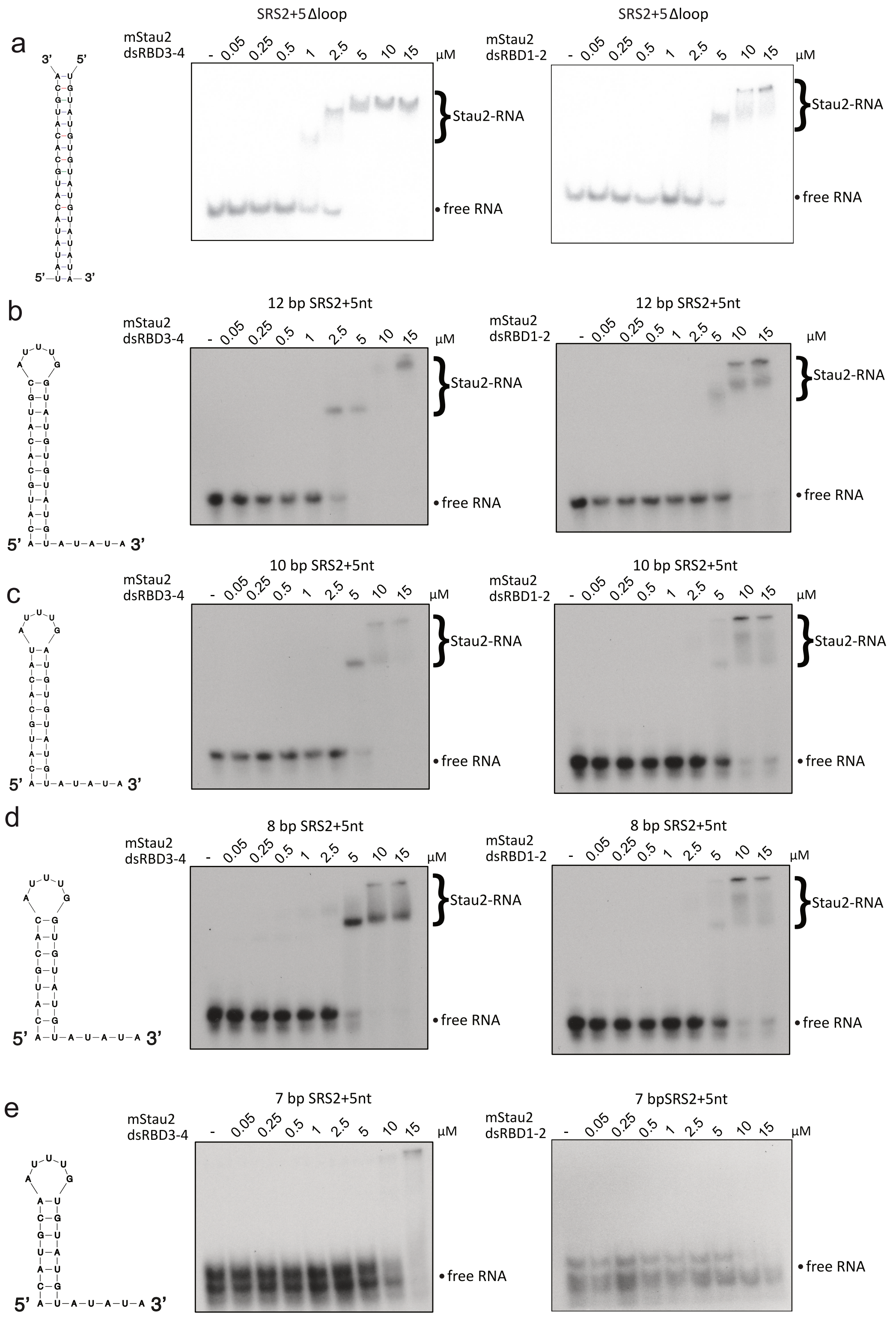


**Supplementary Figure 5: EMSAs with mStau2 tandem domains dsRBD3-4 and dsRBD1-2 and modified SRS2 RNAs. a** mStau2 binding to the elongated stem of SRS2 RNA without the loop region. Binding by both tandem domains to the elongated stem-loop is improved, the stem RNA is bound with similar affinity as SRS2, indicating that total length of the RNA determines binding rather than its specific structure. **b** dsRNA stem-loops with 12bp, **c** 10 bp and **d** 8 bp are bound by both dsRBD3-4 (left) and dsRBD1-2 (right) with similar affinities in the micromolar concentration range. **e** For a stem-loop with only 7 bp, binding is almost completely abolished. No protein-RNA complex is observed. To increase any effects of shortening the stem, SRS2 RNAs with a single-stranded 3' extension were used. This should improve affinity enough to allow for visualization of a deterioration of binding by mStau2 with shortening of the dsRNA stem. Complexes were resolved by native PAGE and imaged by PhosphorImaging or by exposure of radiograph films. Source data are provided as a Source Data file.

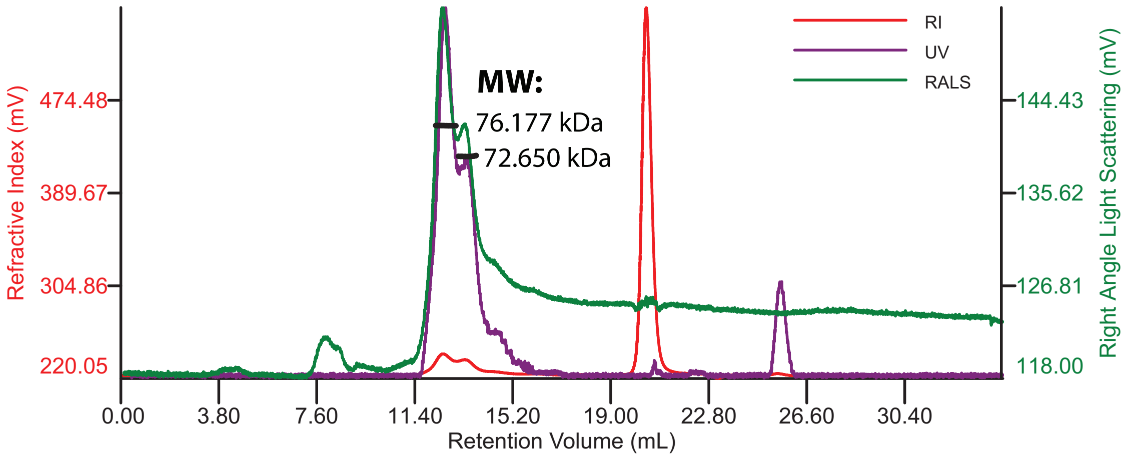


**Supplementary Figure 6: SLS measurements of HisSUMO-Stau2 FL after SEC on 10/300 Superdex200 Increase GL.** SEC chromatogram monitored by UV, refractive index (RI) and right-angle light scattering (RALS). Molecular weight (MW) distribution over peaks 1 and 2 and calculated average MW. The protein elutes in two peaks with molecular weights of 76 kDa and 73 kDa, corresponding to a monomer. The theoretical molecular weight of HisSUMO-tagged Stau2 FL is 75 kDa. The protein co-purified with a slightly smaller degradation product corresponding to peak 2.

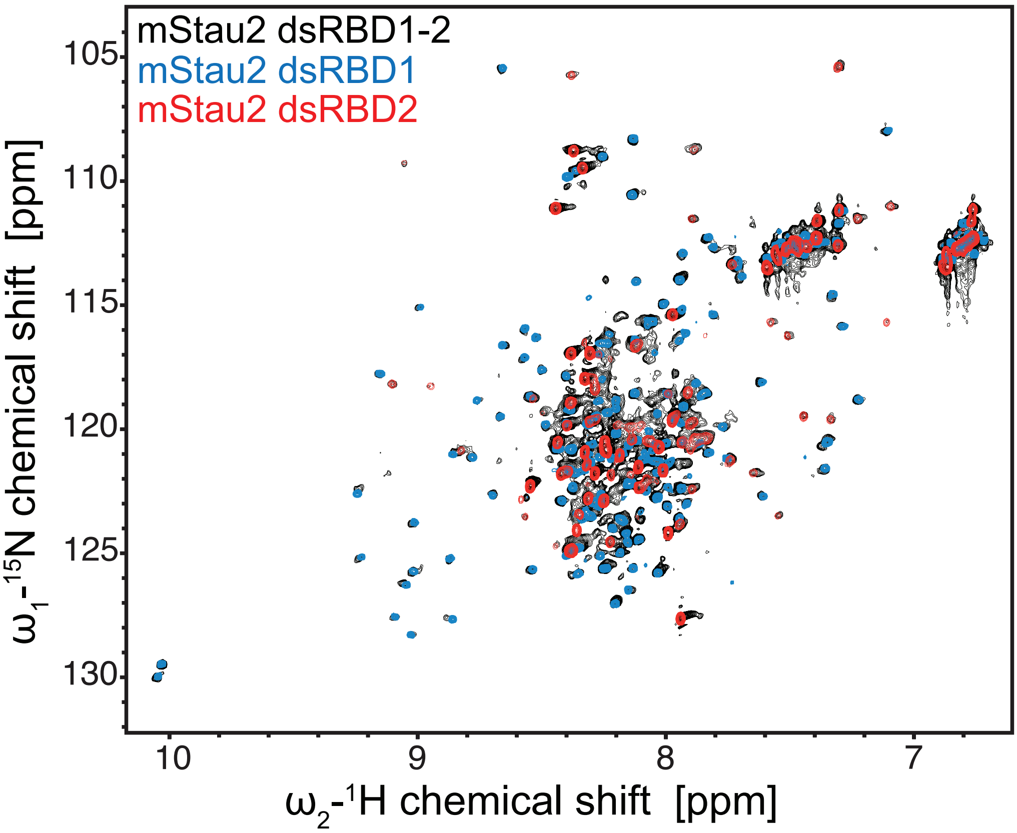


**Supplementary Figure 7: Overlay of the $^{1}H$,$^{15}N$-HSQC spectra of the tandem domain dsRBD1-2 and the individual dsRBDs 1 and 2.** The spectra of dsRBDs 1 and 2 overlap and add up to the spectrum of the tandem domain, indicating that dsRBDs 1 and 2 are separate, partially disordered domains.

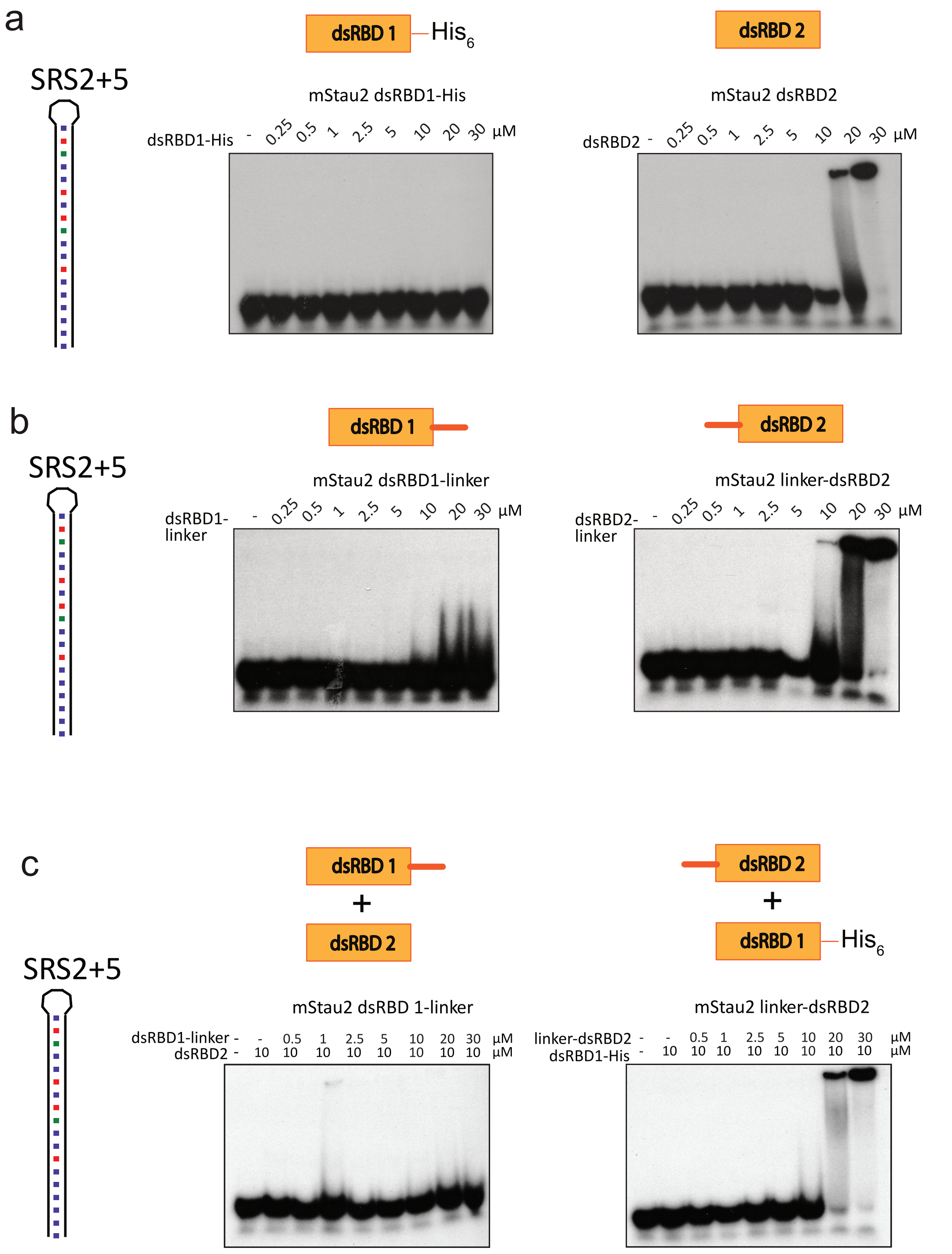


**Supplementary Figure 8: EMSAs with the individual domains dsRBD1 and 2. a** dsSRS2+5 RNA binding of dsRBD1-containing (left) and dsRBD2-containing (right) fragments. **b** Binding of dsRBD1-linker (left) and linker-dsRBD2 (right) to dsSRS2+5 RNA. **c** Binding of dsRBD1-linker (left) and linker-dsRBD2 (right) to dsSRS2+5 RNA in presence of the respective other domain at 10 µM. Source data are provided as a Source Data file.

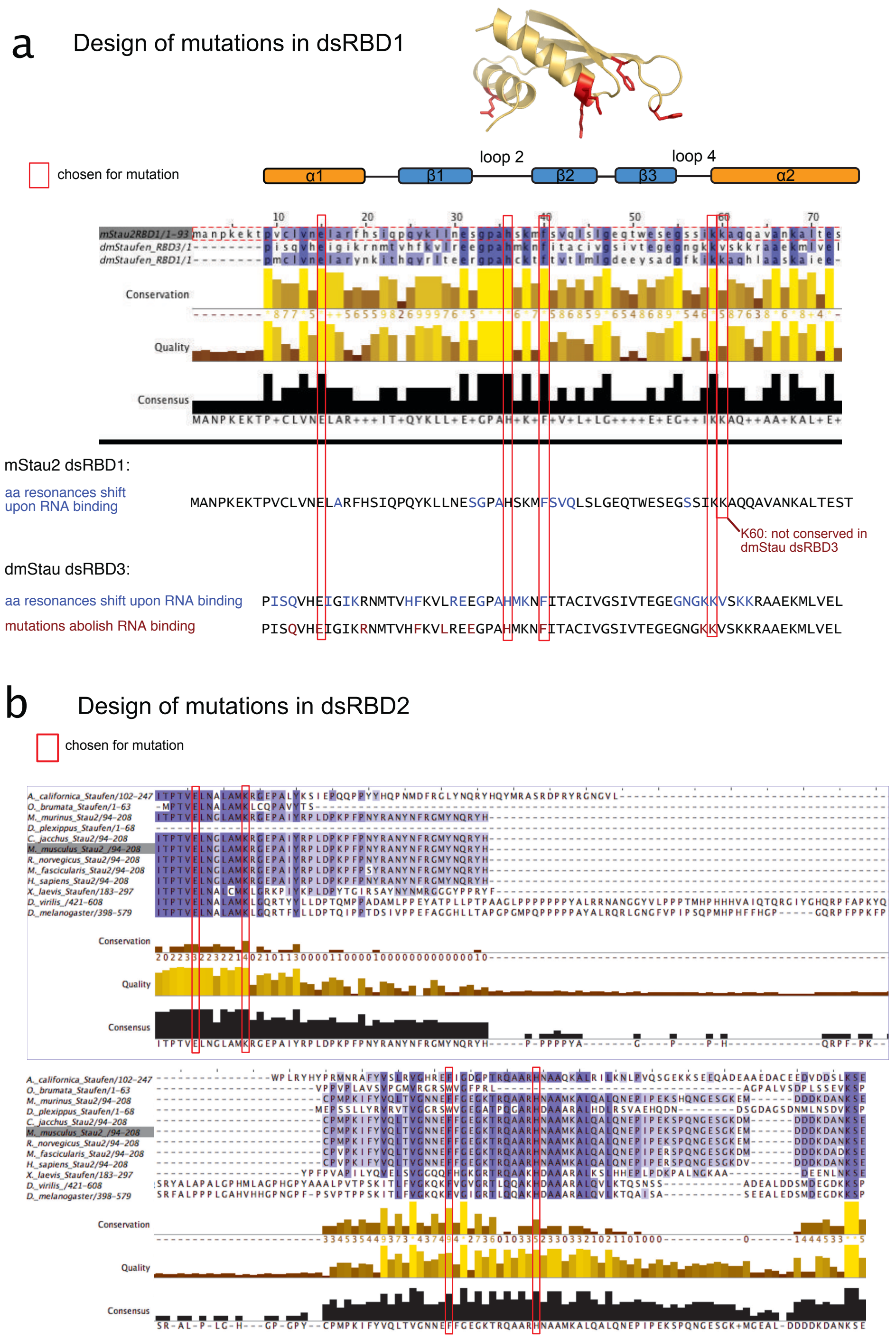


**Supplementary Figure 9: Sequence analysis used for mutant design of dsRBD1 and dsRBD2.** **a** Multiple sequence alignment of mStau2 dsRBD1, dmStau dsRBD1, and dmStau dsRBD3. Below, assigned residues with NMR chemical shift perturbations upon RNA titration are marked in the sequence of mStau2 dsRBD1 in blue. Residues chosen for mutation are marked by red boxes and mapped onto a homology model of dsRBD1. Designed mutations map to regions previously identified to be involved in RNA binding in dmStau dsRBD3[2]. **b** Multiple sequence alignment of mStau2 dsRBD2 to Stau proteins from 11 different species. Residues chosen for mutation are marked by red boxes and mapped onto a homology model of dsRBD2. Plots were generated with the program JalView. Homology models were made using the Phyre2 server[3].

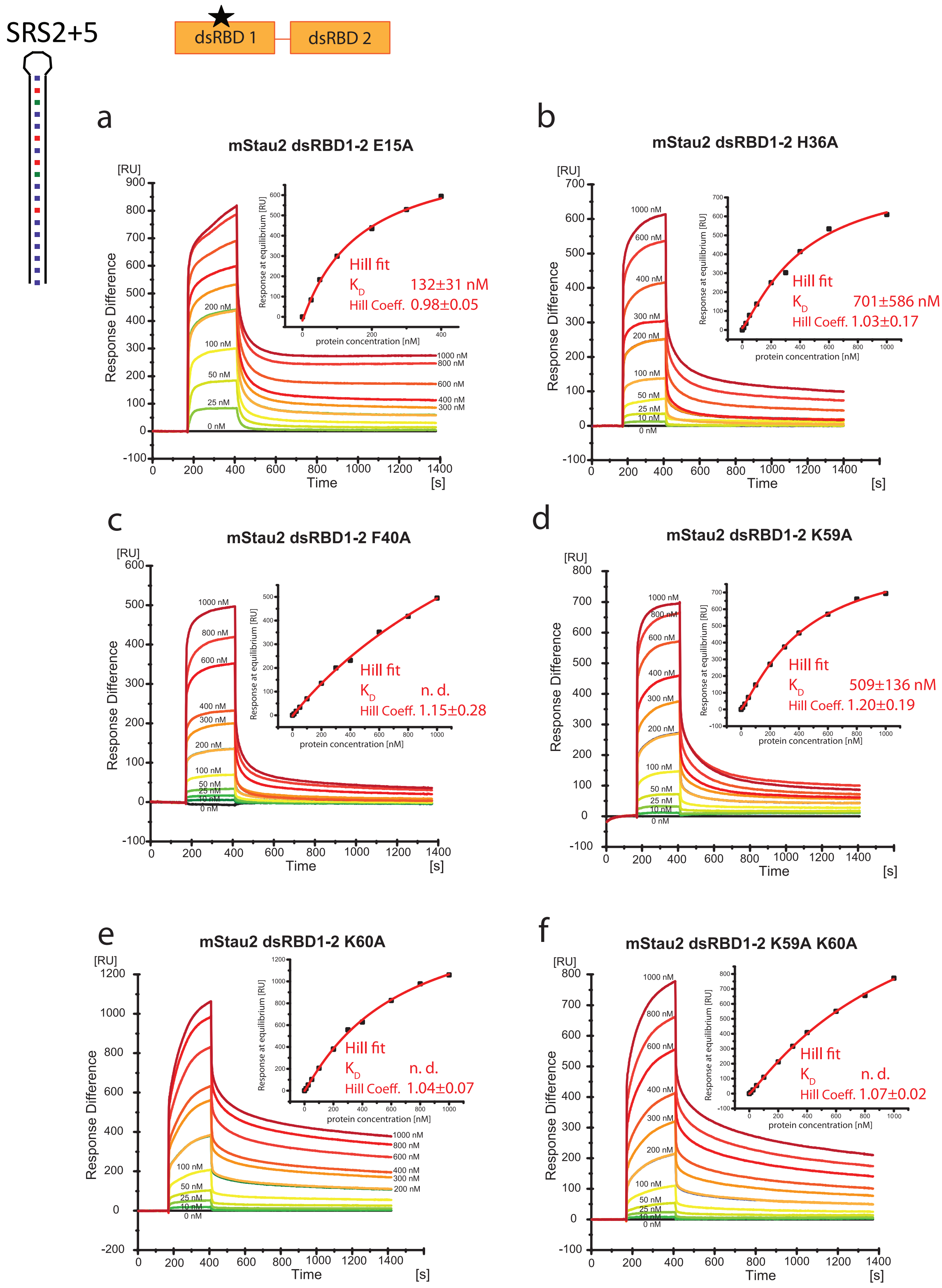


**Supplementary Figure 10: SPR sensorgrams and and curve fitting of SRS2+5 RNA binding by mStau2 dsRBD1-2 with mutations in dsRBD1. a-f** Mutant versions of mStau2 dsRBD1-2. a) E15A, b) H36A, c) F40A, d) K59A, e) K60A and f) K29A K60A bind SRS2+5 transiently with fast kinetics. The steady-state binding curves are described by Hill-fits with Hill coefficients n ≈ 1, indicating non-cooperative binding. Source data are provided as a Source Data file.

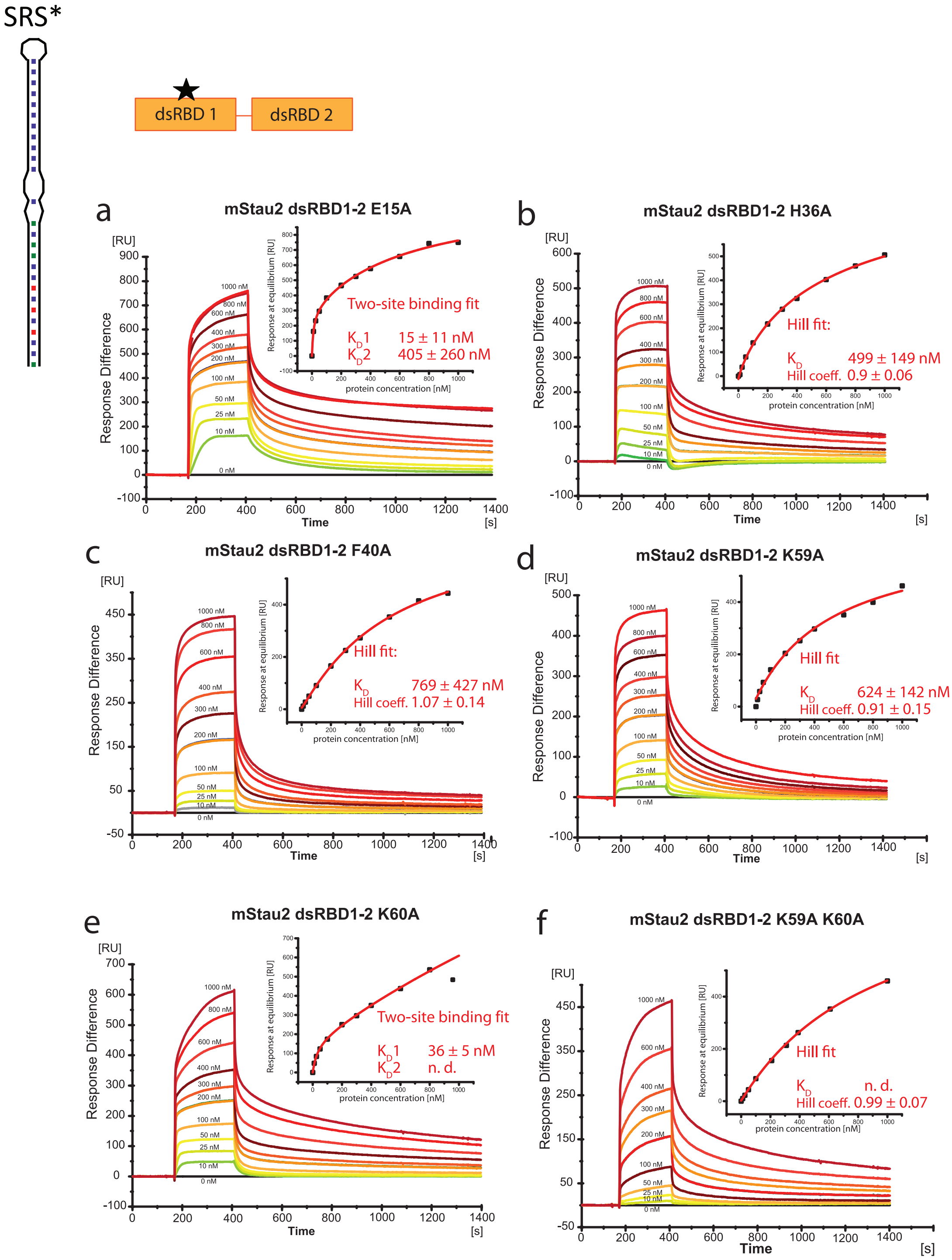


**Supplementary Figure 11: SPR sensorgrams and and curve fitting of SRS* RNA binding by mStau2 dsRBD1-2 with mutations in dsRBD1. a-f** Mutant versions of mStau2 dsRBD1-2. b) H36A, c) F40A, d) K59A and f) K59A K60A bind SRS2+5 transiently with fast kinetics. The steady-state binding curves are described by Hill-fits with Hill coefficients n ≈ 1, indicating non-cooperative binding. Mutants mStau2 dsRBD1-2 a) E15A and e) K60A bind similar to mStau2 dsRBD1-2 wild-type. Source data are provided as a Source Data file.

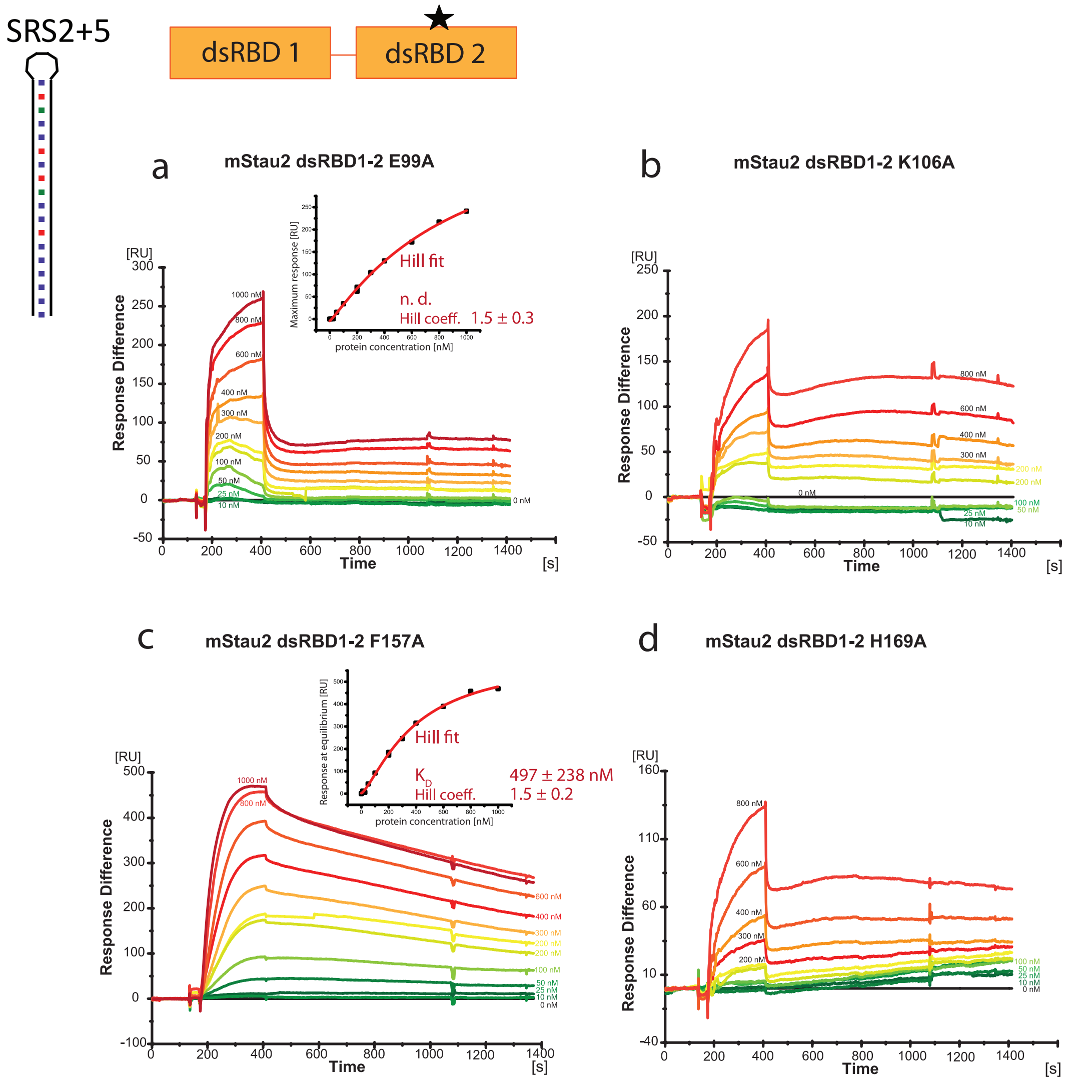


**Supplementary Figure 12: SPR sensorgrams and and curve fitting of SRS2+5 RNA binding by mStau2 dsRBD1-2 with mutations in dsRBD2.** mStau2 dsRBD1-2 **a** E99A, **b** K106A, **c** F157A, **d** H169A show decreased binding to SRS2+5 RNA when compared to wild-type protein. For RBD1-2 E99A (a) and F157A (c), steady-state binding curves are shown. For dsRBD1-2 K106A (b) and H169A (d), steady-state could not be reached. Source data are provided as a Source Data file.

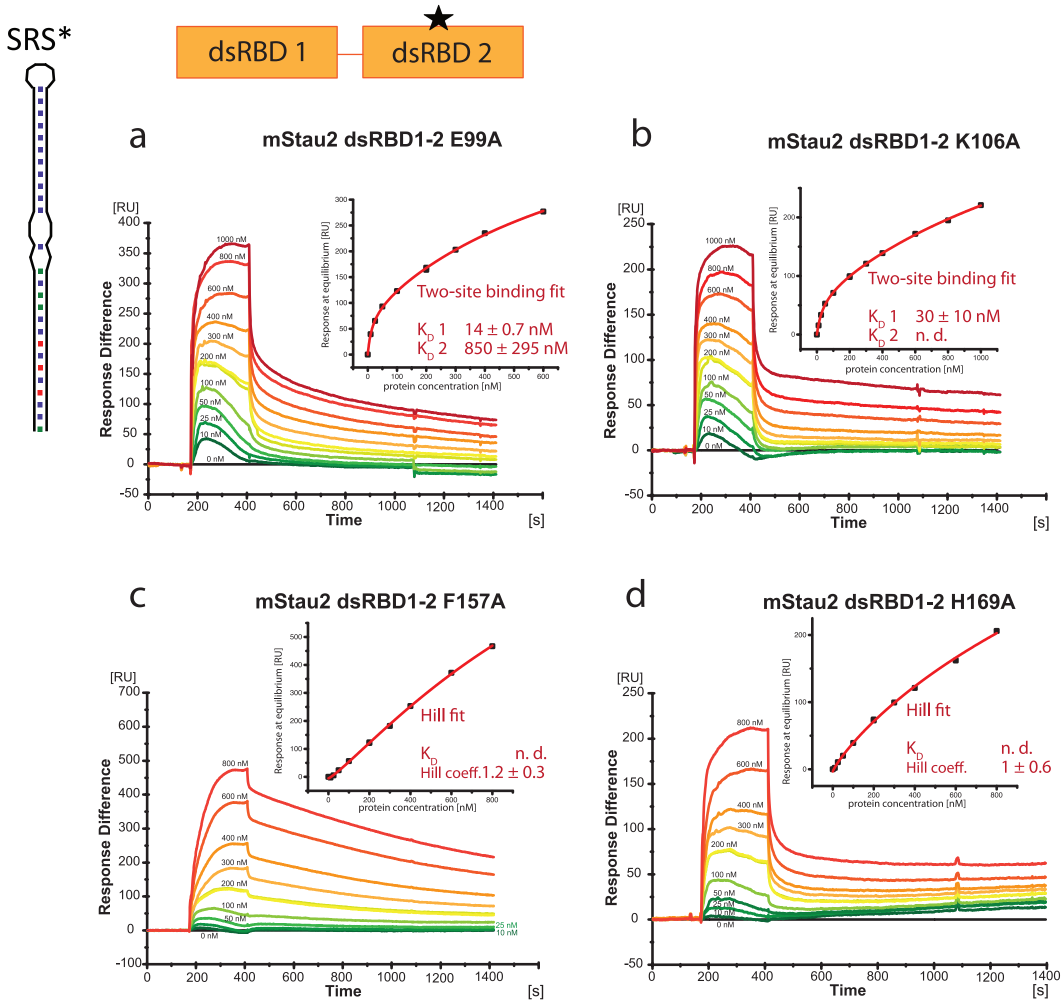


**Supplementary Figure 13: SPR sensorgrams and and curve fitting of SRS* RNA binding by mStau2 dsRBD1-2 with mutations in dsRBD2.** mStau2 dsRBD1-2 **a** E99A, and **b** K106A bind RNA similar to wild-type dsRBD1-2. In contrast, mStau2 dsRBD1-2 **c** F157A and **d** H169A show decreased binding to SRS* RNA. Source data are provided as a Source Data file.

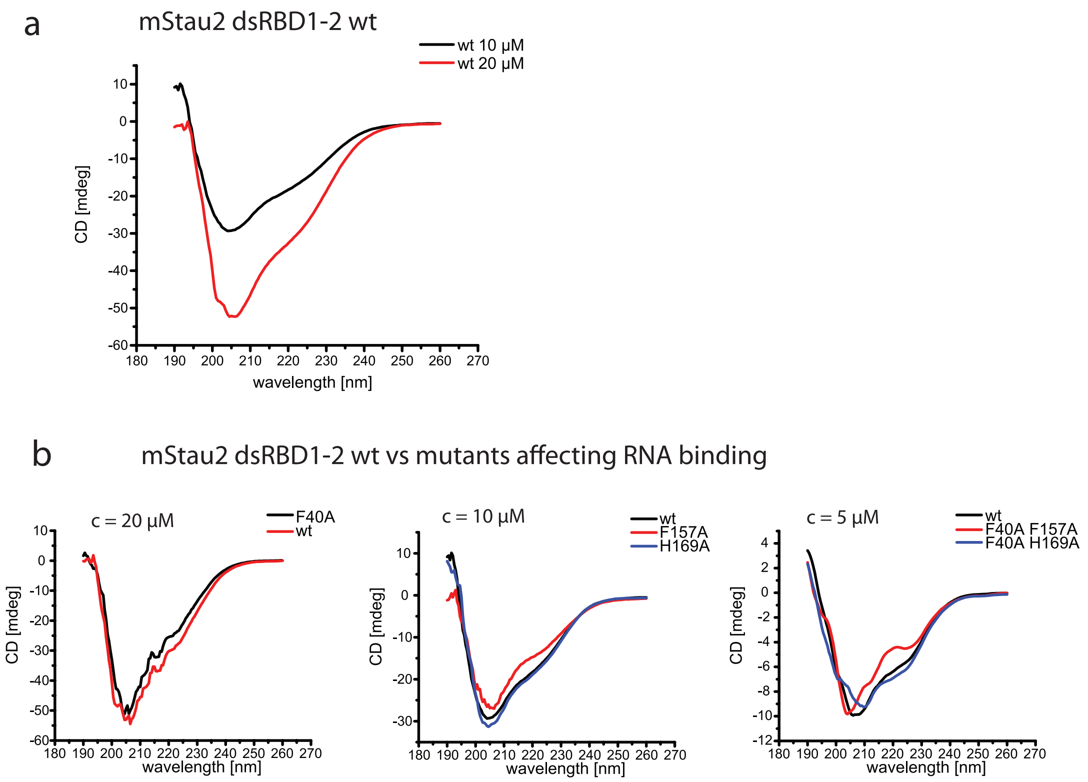


**Supplementary Figure 14: Circular dichroism spectra of mStau2 dsRBD1-2 versions. a** Wild-type mStau2 dsRBD1-2 at different concentrations and **b** RNA-binding mutants of dsRBD1-2 compared to wild-type dsRBD1-2. Measurements were performed at the indicated concentrations in buffer containing <50 mM NaCl. All curves show very similar profiles, indicating that mutant proteins adopt the same fold as wild type dsRBD1-2.

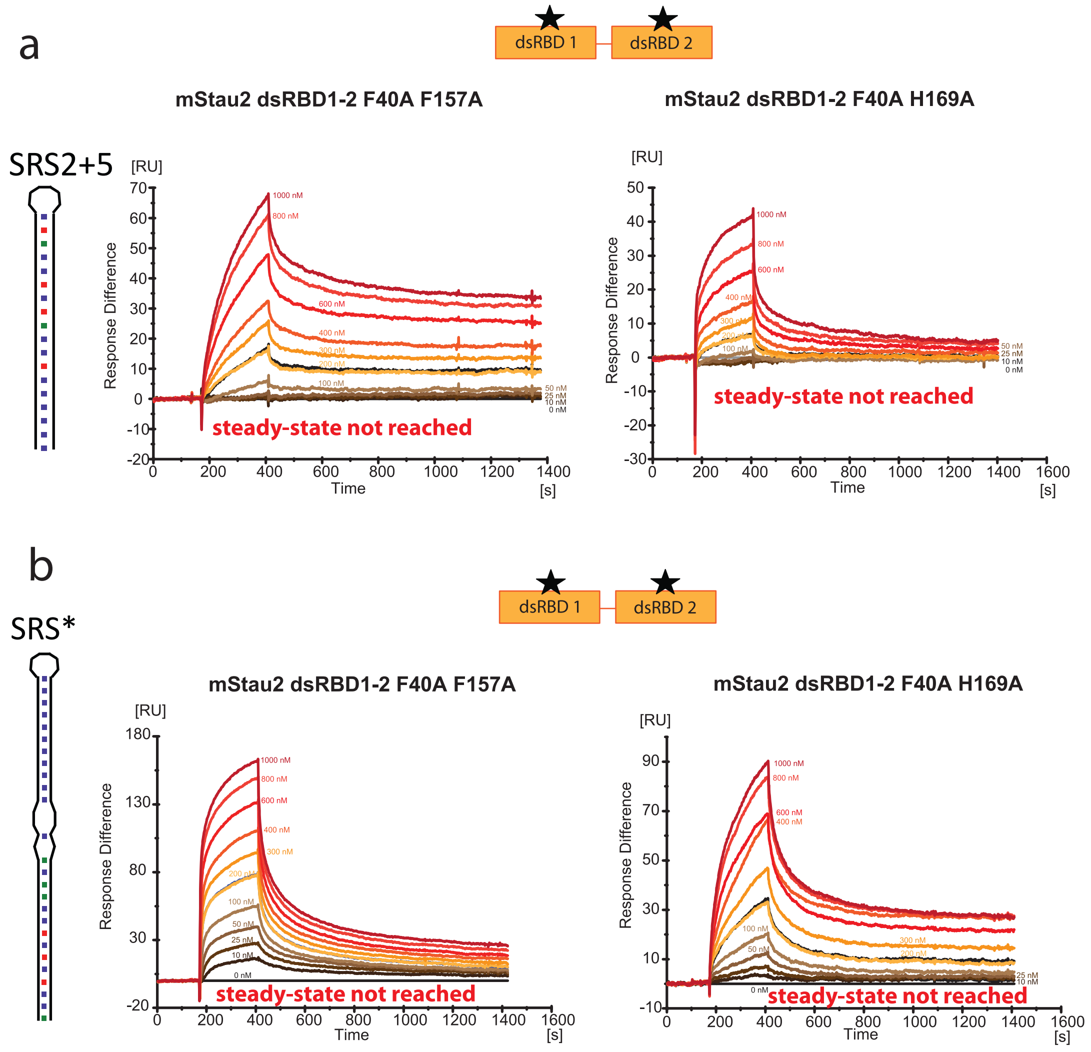


**Supplementary Figure 15: SPR sensorgrams and plots of maximum response against protein concentration of mStau2 dsRBD1-2 double-mutants binding to SRS RNA.** Binding to **a** SRS2+5 and **b** SRS* is strongly decreased as compared to wild-type mStau2 dsRBD1-2. Source data are provided as a Source Data file.

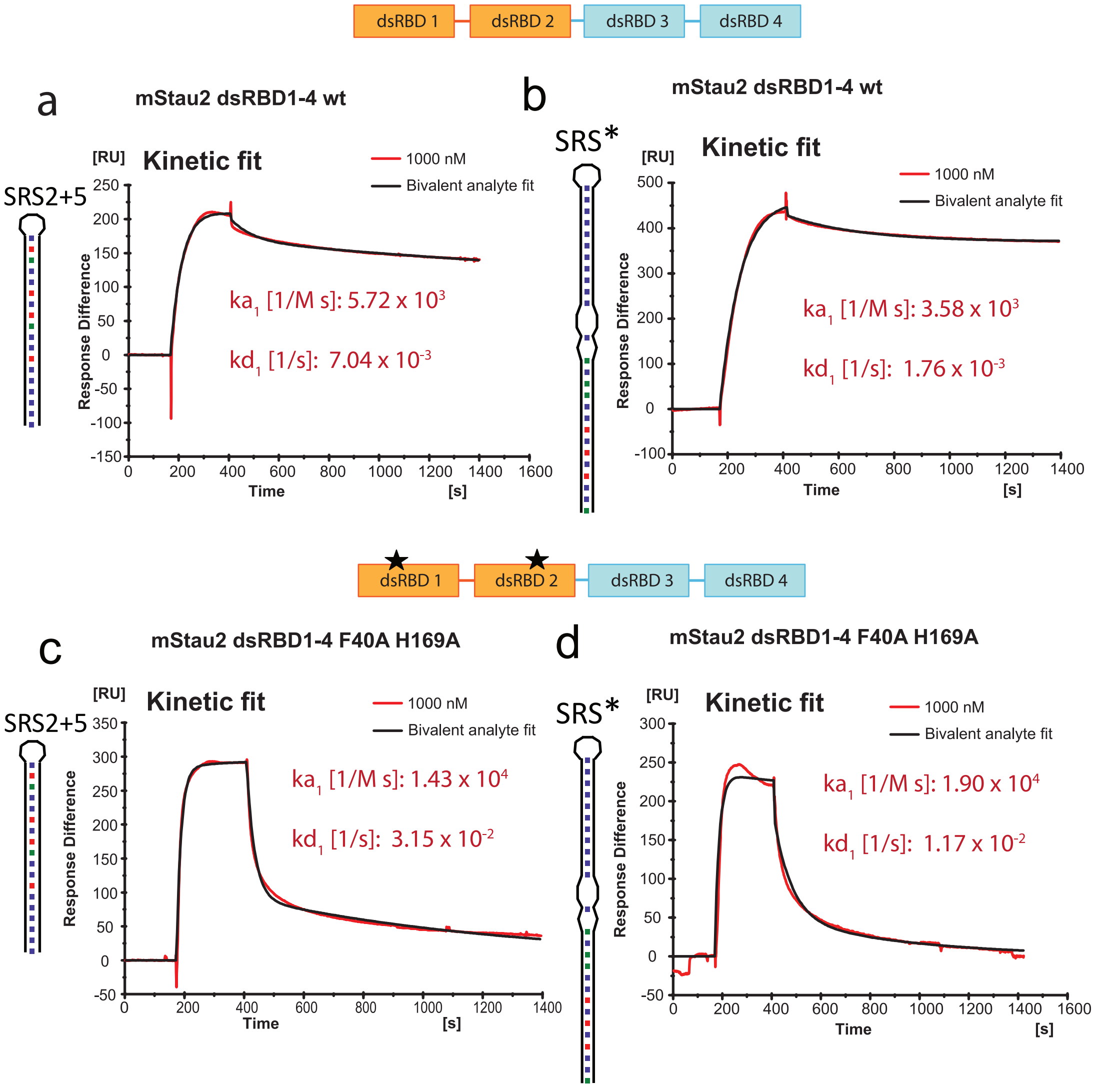


**Supplementary Figure 16: SPR results for mStau2 RBD1-4 and double-mutants binding to RNA.** Exemplary kinetic fits (bivalent analyte fit) at 1000 nM protein concentration are shown. **a** Wild-type mStau2 RBD1-4 binding to SRS2+5 RNA. **b** mStau2 RBD1-4 F40A H169A binding to SRS2+5. The kinetic fit shows that both $ka_1$ and $kd_1$ are ~10-fold increased as compared to wild-type, indicating a transient binding. **c** Wild-type mStau2 RBD1-4 binding to SRS* RNA. **d** mStau2 RBD1-4 F40A H169A binding to SRS*. The kinetic fit shows that both $ka_1$ and $kd_1$ are ~5 and ~10-fold increased, respectively, as compared to wild-type.

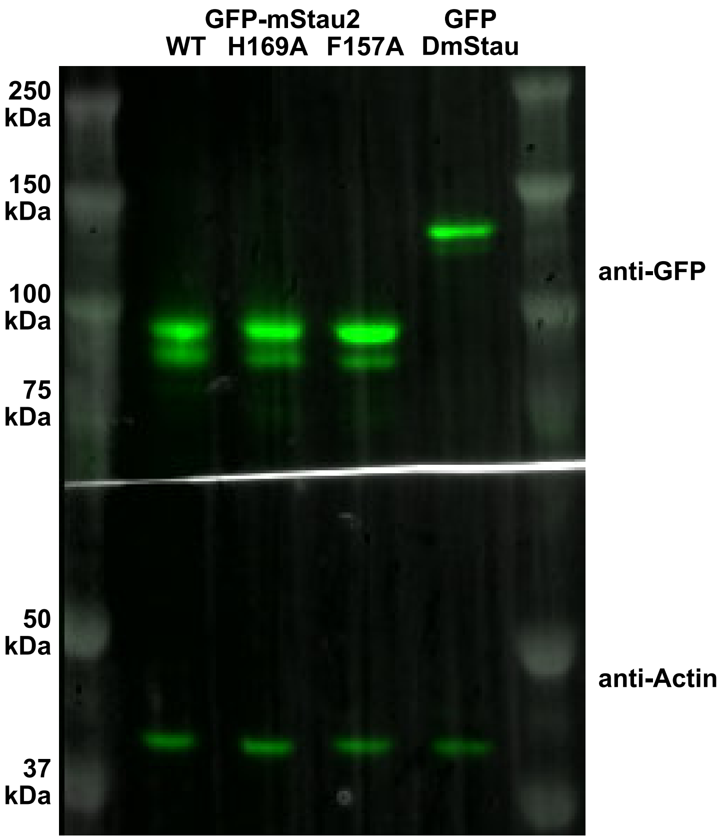


**Supplementary Figure 17**: **wt and mutant Stau proteins are equally expressed in the germline of *stau*$^{R9}$*/stau*$^{D3}$ mutant flies.** Expression levels were checked by Western blot. The blot was developed with anti-GFP (Torrey Pines lab, #TP401) and anti-Actin (Sigma A-2066) primary antibodies and HRP conjugated anti-rabbit secondary antibodies (JacksonImmunoResearch).

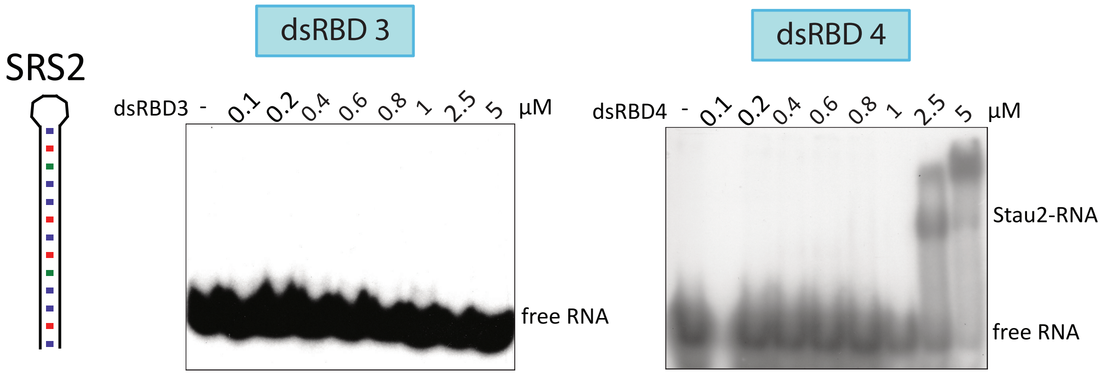


**Supplementary Figure 18: EMSAs with the individual domains dsRBD3 and 4 and SRS2 RNA.** While dsRBD4 binds SRS2 RNA at micromolar concentrations, no binding can be observed for dsRBD3 up to 5 µM protein concentration.

### Supplementary Table 1: Plasmids

| Short name | Full name/description | Source/Cloning strategy |
|---|---|---|
| **pRgs4** | pExpress1<br>- Rgs4 ( (B) ...<br>IMAGp998E0615376Q) | Kiebler lab (LMU München) |
| **pRgs4 3'UTR ¤SRS2** | pEGFP-C2-Rgs4 3'UTR SRS2 deletion 3'UTR | Kiebler lab (LMU München) |
| **pRgs4 3'UTR ¤SRS1** | pEGFP-C2-Rgs4 3'UTR SRS1 deletion | Kiebler lab (LMU München) |
| **pRgs4 3'UTR ¤SRS1 ¤SRS2** | pEGFP-C2-Rgs4 3'UTR SRS1 and SRS2 deletion 3'UTR | Kiebler lab (LMU München) |
| **pET-dmStaufen** | pET3a-Staufen cDNAE10 | Ephrussi lab (EMBL Heidelberg) |
| **prsEGFP2** | | Ephrussi lab (EMBL Heidelberg) |
| **Plasmids created in this study** | | |
| **SH17** | pCR-II-Blunt-TOPO-Rgs4 BR | Blunt end TOPO cloning |
| **SH22** | pOPINS3C-mStau2 FL<br>(Staufen homolog 2 isoform 3 [Mus Musculus],<br>NCBI Reference Sequence:<br>NP_079579.2) | InFusion cloning,<br>primers Stau2 FW and Stau2 RV |
| **SH23** | pOPINS3C-mStau2 RBD3-4<br>(mStau2 200-373) | InFusion cloning,<br>primers RBD3 FW and RBD4 RV |
| **SH27** | pOPINS3C-Stau2 RBD4<br>(mStau2 272-373) | InFusion cloning,<br>primers RBD4 FW and RBD4 RV |
| **SH29** | pOPINS3C-mStau2 RBD1-4<br>(mStau2 1-373) | InFusion cloning,<br>primers Stau2 FW and RBD4 RV |
| **SH30** | pFastBacDual-HisSUMO-mStau2 FL<br>(Staufen homolog 2 isoform 3 [Mus Musculus],<br>NCBI Reference Sequence:<br>NP_079579.2) | InFusion cloning, primers His-SUMO FW and mStau2 RV-pFBD |
| **SH31** | pOPINS3C-mStau2 RBD1-2<br>(mStau2 1-208) | InFusion cloning,<br>primers Stau2 FW and RBD2 RV |
| **SH41** | pOPINS3C-mStau2 RBD1-2 E15A<br>(mStau2 1-208 E15A) | InFusion cloning,<br>3-point PCR with primers Stau2 FW+Stau2 E15A antisense, Stau2 E15A sense+RBD2 RV |
| **SH42** | pOPINS3C-mStau2 RBD1-2 K59A K60A<br>(mStau2 1-208 K59A K60A) | InFusion cloning,<br>3-point PCR with primers Stau2 FW+Stau2 K59A K60A antisense, Stau2 K59A K60A sense+RBD2 RV |
| **SH45** | pOPINS3C-mStau2 RBD2<br>(mStau2 94-208) | InFusion cloning,<br>primers RBD2 SM FW and RBD2 RV |
| **SH46** | pOPINJ-mStau2 RBD1-6xHis<br>(mStau2 (1-74) - 6xHis) | InFusion cloning,<br>PCR1 with Stau2 FW and RBD1+6xHis RV, PCR2 on product of PCR1 with Stau2 FW and pOPIN-6xHis RV |
| **SH47** | pOPINS3C-mStau2 RBD1 linker<br>(mStau2 1-93) | InFusion cloning,<br>primers Stau2 FW and RBD1-linker RV |
| **SH48** | pOPINS3C-mStau2 linker-RBD2<br>(mStau2 75-208) | InFusion cloning,<br>primers linker-RBD2 FW and RBD2 RV |
| **SH49** | pOPINS3C-mStau2 RBD1-2 H36A<br>(mStau2 1-208 H36A) | InFusion cloning,<br>3-point PCR with primers Stau2 FW+Stau2 H36A antisense, Stau2 H36A sense+RBD2 RV |

| Short name | Full name/description | Source/Cloning strategy |
|---|---|---|
| **SH50** | pOPINS3C-mStau2 RBD1-2 F40A (mStau2 1-208 F40A) | InFusion cloning, 3-point PCR with primers Stau2 FW+Stau2 F40A antisense, Stau2 F40A sense+RBD2 RV |
| **SH51** | pOPINS3C-mStau2 RBD1-2 K59A (mStau2 1-208 K59A) | InFusion cloning, 3-point PCR with primers Stau2 FW+Stau2 K59A antisense, Stau2 K59A sense+RBD2 RV |
| **SH52** | pOPINS3C-mStau2 RBD1-2 K60A (mStau2 1-208 K60A) | InFusion cloning, 3-point PCR with primers Stau2 FW+Stau2 K60A antisense, Stau2 K60A sense+RBD2 RV |
| **SH53** | pBlueScript-KS-rsEGFP2-mStau2 FL | InFusion cloning, restriction enzymes BamHI and XbaI, 3-point PCR with primers pBSKS-rsEGFP2 FW + rsEGFP+3C RV, 3C+Stau2 FW+ pBSKS-Stau2 RV |
| **SH55** | pOPINS3C-mStau2 RBD1-2 E99A (mStau2 1-208 E99A) | InFusion cloning, 3-point PCR with primers Stau2 FW+Stau2 E99A antisense, Stau2 E99A sense+RBD2 RV |
| **SH56** | pOPINS3C-mStau2 RBD1-2 F157A (mStau2 1-208 F157A) | InFusion cloning, 3-point PCR with primers Stau2 FW+Stau2 F1570A antisense, Stau2 F157A sense+RBD2 RV |
| **SH58** | pOPINS3C-mStau2 RBD1-2 H169A (mStau2 1-208 H169A) | InFusion cloning, 3-point PCR with primers Stau2 FW+Stau2 H169A antisense, Stau2 H169A sense+RBD2 RV |
| **SH59** | pUASp attb-rsEGFP2-mStau2 | Infusion cloning with primers pUASp-rsEGFP2 FW and pUASp-Stau2 RV, template SH53 |
| **SH60** | pOPINS3C-mStau2 RBD1-2 K106A (mStau2 1-208 K106A) | InFusion cloning, 3-point PCR with primers Stau2 FW+Stau2 K106A antisense, Stau2 K106A sense+RBD2 RV |
| **SH64** | pOPINS3C-mStau2 RBD1-2 F40A H169A (mStau2 1-208 F40A H169A) | InFusion cloning, 3-point PCR with primers Stau2 FW+Stau2 F40A antisense, Stau2 F40A sense+RBD2 RV on SH58 as template |
| **SH65** | pOPINS3C-mStau2 RBD1-2 F40A H169A (mStau2 1-208 F40A F157A) | InFusion cloning, 3-point PCR with primers Stau2 FW+Stau2 F40A antisense, Stau2 F40A sense+RBD2 RV on SH56 as template |
| **SH66** | pBlueScript-KS-rsEGFP2-mStau2 F40A | Site-directed mutagenesis (QuikChange II kit) with Stau2 F40A sense and Stau2 F40A antisense |
| **SH67** | pBlueScript-KS-rsEGFP2-mStau2 F40A F157A | Site-directed mutagenesis (QuikChange II kit) on SH66 with Stau2 F157A sense and Stau2 F157A antisense |
| **SH68** | pBlueScript-KS-rsEGFP2-mStau2 F40A H169A | Site-directed mutagenesis (QuikChange II kit) on SH66 with Stau2 H169A sense and Stau2 H169A antisense |
| **SH69** | pOPINS3C-mStau2 RBD1-4 F40A H169A | InFusion cloning with primers Stau2 FW and RBD4 RV on template SH68 |

| Short name | Full name/description | Source/Cloning strategy |
|---|---|---|
| **SH70** | pOPINS3C-mStau2 RBD1-4 F40A F157A | InFusion cloning with primers Stau2 FW and RBD4 RV on template SH67 |
| **SH71** | pUASp attB-rsEGFP2-mStau2 F40A F157A | Infusion cloning with primers pUASp-rsEGFP2 FW and pUASp-Stau2 RV, template SH67 |
| **SH72** | pUASp attB-rsEGFP2-mStau2 F40A H169A | Infusion cloning with primers pUASp-rsEGFP2 FW and pUASp-Stau2 RV, template SH68 |

### Supplementary Table 2: Cloning primers

| Primer name | Primer description | Sequence 5' → 3' |
|---|---|---|
| **mStau2 FW** | pOPIN- mStau2 FW +PPsite | AAGTTCTGTTTCAGGGCCCGATGGCAAACCCCAAAGAGA |
| **RBD2 FW** | pOPIN- mStau2 RBD2 FW +PPsite | AAGTTCTGTTTCAGGGCCCGATGCCCAAGATCTTTTATGTTCAGT |
| **Linker-RBD2 FW** | pOPIN- mStau2 RBD2 long FW +PPsite | AAGTTCTGTTTCAGGGCCCGCTTCCCAAACCAGTTCAGAAAC |
| **RBD3 FW** | pOPIN- mStau2 RBD3 FW +PPsite | AAGTTCTGTTTCAGGGCCCGATAAGCTTAGTGTTTGAGATTGCGC |
| **RBD1-linker RV** | mStau2 RBD1-linker RV-pOPIN | CTGGTCTAGAAAGCTTCTCTAACTACCTGGGTTATTATTGACATTAC |
| **RBD2 RV** | mStau2 RBD2 RV-pOPIN | CTGGTCTAGAAAGCTTCTATTCAGATTTATTTGCATCTTTATCGTC |
| **RBD4 RV** | mStau2 RBD4 RV-pOPIN | CTGGTCTAGAAAGCTTCTAAAGCTGTAACAGCATTGCTT |
| **Stau2 RV** | mStau2 FL RV-pOPIN | CTGGTCTAGAAAGCTTCTAGATGGCCGACTTTGAT |
| **pOPIN-6xHis RV** | pOPIN-6xHis RV | CTGGTCTAGAAAGCTTCTAgtgatggtggtgatggtg |
| **RBD1-6xHis RV** | RBD1-6xHis RV | gtgatggtggtgatggtgCGTAGATTCGTCAAACGCTT |
| **His-SUMO FW** | pFastBacDual-His-SUMO FW | CATCGGGCGCGGATCCATGGCACACCATCACCAC |
| **Stau2 FL RV-pFBD** | Stau2 FL RV-pFastBacDual | ACTTCTCGACAAGCTTCTAGATGGCCGACTTTGAT |
| **pBSKS-rsEGFP2 FW** | pBSKS-rsEGFP2 FW | gcggtggcggccgctctagaatggtgagcaagggcga |
| **rsEGFP+3C RV** | rsEGFP+3C RV | cgggccctgaaacagaacttccagcttgtacagctcgtccatgc |
| **3C+mStau2 FW** | 3C+mStau2 FW | ctggaagttctgtttcagggcccgATGGCAAACCCCAAAGAGAA |
| **pBSKS-mStau2 RV** | pBSKS-mStau2 RV | tcctgcagcccgggggatccCTAGATGGCCGACTTTGATTTCT |
| **pUASp-rsEGFP2 FW** | pUASp-rsEGFP2 FW | AGGCCACTAGTGGATCTGGATCCatggtgagcaagggcga |
| **pUASp-mStau2 RV** | pUASp-mStau2 RV | TTAACGTTCGAGGTCGACTCTAGACTAGATGGCCGACTTTGATT |

### Supplementary Table 3: Mutagenesis primers

| Primer name | Primer description | Sequence 5’ → 3’ |
|---|---|---|
| **mStau2 E15A antisense** | a44c_antisense | ggaaacgggctaacgcatttaccagacacactggag |
| **Stau2 E15A sense** | a44c | ctccagtgtgtctggtaaatgcgttagcccgtttcc |
| **mStau2 H36A antisense** | c106g_a107c antisense | caccgaaaacatcttcgaagcagcaggcccgctttcattc |
| **mStau2 H36A sense** | c106g_a107c_ | gaatgaaagcgggcctgctgcttcgaagatgttttcggtg |
| **mStau2 F40A antisense** | t118g_t119c_ antisense | agactcagctgcaccgaagccatcttcgaatgagcagg |
| **mStau2 F40A sense** | t118g_t119c | cctgctcattcgaagatggcttcggtgcagctgagtct |
| **mStau2 K59A antisense** | a175g_a176c_ antisense | ttgttgggccttcgctatactgctcccttcggattccc |
| **Stau2 K59A sense** | a175g_a176c | gggaatccgaagggagcagtatagcgaaggcccaaca a |
| **mStau2 K60A antisense** | a178g_a179c_antis ense | caacagcttgttgggccgcctttatactgctcccttcgg |
| **Stau2 K60A sense** | a178g_a179c_ | ccgaagggagcagtataaaggcggcccaacaagctgtt g |
| **mStau2 K59A K60A antisense** | a175g_a176c_a178 g_a179c_antisense | cagcttgttgggccgccgctatactgctcccttcggattccca tgt |
| **mStau2 K59A K60A sense** | a175g_a176c_a178 g_a179c | acatgggaatccgaagggagcagtatagcggcggccca acaagctg |
| **mStau2 ¤119-135 antisense** | | ggcaatgatacctctgtggatctagtggcctg |
| **mStau2 ¤119-135 sense** | | caggccactagatccacagaggtatcattgcc |
| **mStau2 E99A antisense** | a296c_antisense | tagcgagcccattcagtgccacagttggagttata |
| **Stau2 E99A sense** | a296c_sense | tataactccaactgtggcactgaatgggctcgcta |
| **mStau2 K106A antisense** | a316g_a317c antisense | ggcaggctctccccttgccatagcgagcccattc |
| **mStau2 K106A sense** | a316g_a317c | gaatgggctcgctatggcaaggggagagcctgcc |
| **mStau2 F157A sense** | t469g_t470c | gttcagttaactgtaggaaataatgaagcctttggtgaagg gaagactc |
| **mStau2 F157A antisense** | t469g_t470c antisense | gagtcttcccttcaccaaaggcttcattatttcctacagttaac tgaac |
| **mStau2 H169A antisense** | c505g_a506c antisense | ctttcatcgcagcattggctctggcagcttgtcgag |
| **mStau2 H169A sense** | c505g_a506c | ctcgacaagctgccagagccaatgctgcgatgaaag |

### Supplementary Table 4: Template primers for RNA *in vitro* transcription

| Primer name | Primer description | Sequence 5’ → 3’ |
|---|---|---|
| **3’UTR FW** | T7prom+*Rgs4* 3’UTR FW | AATTTAATACGACTCACTATAGGttctc acacagaggcagagaacc |
| **3’UTR RV** | *Rgs4* 3’UTR 2129 RV | aggcctataaagcacatggcagaaacagacat |
| **BR FW** | T7prom+*Rgs4* 3’UTR 257FW | AATTTAATACGACTCACTATAGGtaat ggccctgtaggtctgg |
| **BR RV** | *Rgs4* 3’UTR 870 RV | acgtgagcaaccaaccac |
| **T7 FW** | T7prom FW | AATTTAATACGACTCACTATAGG |
| **SRS1 RV** | SRS1-T7prom RV | GCTCCATCAAGACCCAGTGGCTTGA CGGAACCCTATAGTGAGTCGTATTA AATT |

| Primer name | Primer description | Sequence 5’ → 3’ |
|---|---|---|
| **SRS2 RV** | SRS2-T7prom RV | ACATACACATACACAAATTGCATGT<br>GCATGTCCCTATAGT<br>GAGTCGTATTAAATT |
| **SRS3 RV** | SRS3-T7prom RV | Catgtgtgaacatatacaaatatatatatgcatatata<br>tatatattcatgcacattcacacataCCTATAGT<br>GAGTCGTATTAAATT |
| **SRS2+5 RV** | SRS2+5-T7prom RV | TATATACATACACATACACAAATTGC<br>ATGTGCATGTATATACCTATAGTGA<br>GTCGTATTAAATT |
| **11bpSRS2+5 RV** | 11bpSRS2+5- T7prom RV | TATATACATACACATACAAATCATGT<br>GCATGTCCTATAGTGAGTCGTATTA<br>AATT |
| **9bpSRS2+5 RV** | 9bpSRS2+5-T7prom RV | TATATACATACACACAAATTGTGCAT<br>GTCCTATAGTGAGTCGTATTAAATT |
| **7bpSRS2+5 RV** | 7bpSRS2+5-T7prom RV | TATATACATACACAAATTGCATGTCC<br>TATAGTGAGTCGTATTAAATT |
| **6bpSRS2+5 RV** | 6bpSRS2+5-T7prom RV | TATATACATACCAAATGCATGTCCTA<br>TAGTGAGTCGTATTAAATT |

### Supplementary Table 5: RNA sequences

| Short name | Full name/<br>description<br>(numbering relative to start of 3’UTR) | Production |
|---|---|---|
| ***Rgs4* 3’UTR** | *Rgs4* 3’UTR FL<br><br>Rattus norvegicus regulator of G-protein signaling 4 (Rgs4), mRNA<br>NCBI Reference Sequence: NM_017214.1<br>(3’UTR: nt 728-2919 of mRNA) | *In vitro* transcription<br>(Ambion MegaScript),<br>primer: 3’UTR FW, 3’UTR RV |
| **BR** | *Rgs4* 3’UTR (257-890) | *In vitro* transcription<br>(Ambion MegaScript),<br>primer: BR FW, BR RV |
| **BR ¤SRS2** | *Rgs4* 3’UTR (257-890) Δ(354-366)(417-429) | *In vitro* transcription<br>(Ambion MegaScript),<br>primer: BR FW, BR RV |
| **BR ¤SRS1** | *Rgs4* 3’UTR (257-890)Δ(729-759) | *In vitro* transcription<br>(Ambion MegaScript),<br>primer: BR FW, BR RV |
| **BR ¤SRS1 ¤SRS2** | *Rgs4* 3’UTR (257-890)Δ(354-366)(417-429) (729-759) | *In vitro* transcription<br>(Ambion MegaScript),<br>primer: BR FW, BR RV |
| **SRS1** | *Rgs4* 3’UTR (729-759) | *In vitro* transcription (Ambion MegaShortScript),<br>Chemical synthesis (IBA) |
| **SRS2** | *Rgs4* 3’UTR (354-429)-Δ(367-416)AUUUG | *In vitro* transcription (Ambion MegaShortScript),<br>Chemical synthesis (Dharmacon & IBA) |
| **SRS3** | *Rgs4* 3’UTR (492-555) | *In vitro* transcription (Ambion MegaShortScript) |
| **SRS2+5** | *Rgs4* 3’UTR (349-434)-Δ(367-416)AUUUG | *In vitro* transcription, primers: T7prom, dsSRS2+5 RV |

| Short name | Full name/ description (numbering relative to start of 3’UTR) | Production |
|---|---|---|
| **11bpSRS2+5** | *Rgs4* 3’UTR (354-434)-Δ(369-418)AUUUG | *In vitro* transcription (Ambion MegaShortScript), primers: T7prom, 11bpSRS2+5 RV |
| **9bpSRS2+5** | *Rgs4* 3’UTR (354-434)-Δ(371-419)AUUUG | *In vitro* transcription (Ambion MegaShortScript), primers: T7prom, 9bpSRS2+5 RV |
| **7bpSRS2+5** | *Rgs4* 3’UTR (354-434)-Δ(373-421)AUUUG | *In vitro* transcription (Ambion MegaShortScript), primers: T7prom, 7bpSRS2+5 RV |
| **6bpSRS2+5** | *Rgs4* 3’UTR (354-434)-Δ(374-422)AUUUG | *In vitro* transcription (Ambion MegaShortScript), primers: T7prom, 6bpSRS2+5 RV |

#### Supplementary Table 6: Sequences of ssDNA oligonucleotides specific to *bicoid*

| Name | Sequence |
|---|---|
| **bcd_5UTR_1** | TGGCAAAGGAGTGTGGAAAC |
| **bcd_CD_1** | CTGAAGCTGCGGATGTTGG |
| **bcd_CD_2** | TCGAAGGGATTTCGGAATTG |
| **bcd_CD_3** | CCATATCTTCACCTGGGCTG |
| **bcd_CD_4** | GTCCTTGTGCTGATCCGAT |
| **bcd_CD_5** | CTCCACCCAAGCTAAGAGTC |
| **bcd_CD_6** | GCGTTGAATGACTCGCTGTAG |
| **bcd_CD_7** | TGTGGCCTCCATTGTAGTTG |
| **bcd_CD_8** | GGTGATTATGGACCTGCTGC |
| **bcd_CD_9** | GCTGGAAGTCAAAGTGATGG |
| **bcd_CD_10** | GTAGTACGAGCTGTTGAAGTTG |
| **bcd_CD_11** | GTGTTAATGGCTCGTAGACC |
| **bcd_CD_12** | CACACAGACTCGGACTTTCG |
| **bcd_CD_13** | CTTCTTGCTCGTTCCGTCG |
| **bcd_CD_14** | CCCTTCAAAGGCTCCAAGATC |
| **bcd_CD_15** | CTAAGGCTCTTATTCCGGTGC |
| **bcd_CD_16** | CTCCACGATTTCCGGTTCC |
| **bcd_CD_17** | GCTTGCATTATCGTATCCATCG |
| **bcd_CD_18** | CATCCAGGCTAATTGAAGCAG |
| **bcd_3'UTR_1** | ATGAAACTCTCTAACACGCCTC |
| **bcd_3'UTR_2** | GTACAATCAGGAACAACAGTGG |
| **bcd_3'UTR_3** | ACACGGATCTTAGGACTAGACC |
| **bcd_3'UTR_4** | GAATAGCGTATTGCAGGGAAAG |
| **bcd_3'UTR_5** | GCCCAAATGGCCTCAAATG |
| **bcd_3'UTR_6** | CCGAAATGTGGGACGATAAC |

### Supplementary References

1. Heraud-Farlow JE, *et al.* Staufen2 Regulates Neuronal Target RNAs. *Cell reports* **5**, 1511-1518 (2013).

2. Ramos A, *et al.* RNA recognition by a Staufen double-stranded RNA-binding domain. *EMBO J* **19**, 997-1009 (2000).

3. Kelley LA, Mezulis S, Yates CM, Wass MN, Sternberg MJ. The Phyre2 web portal for protein modeling, prediction and analysis. *Nature protocols* **10**, 845-858 (2015).
